## Supplementary Table 1 for "VirION2: a short- and long-read sequencing and informatics workflow to study the genomic diversity of viruses in nature"

**Supplementary Table 1:** Virus mock community composition and characteristics

| Phage | Taxonomy | GC (%) | Genome Size (kb) |
| --- | --- | --- | --- |
| *Pseudoalteromonas* phage HM1 | *Myoviridae* | 35.7 | 129.4 |
| *Pseudoalteromonas* phage HP1 | *Podoviridae* | 44.7 | 45.0 |
| *Pseudoalteromonas* phage HS2 | *Siphoviridae* | 40.2 | 38.2 |
