## Supplementary Table 2 for "VirION2: a short- and long-read sequencing and informatics workflow to study the genomic diversity of viruses in nature"

**Supplementary Table 2:** Error-correction performance benchmarks between assembly methods and sequencing libraries.

| **Assembly** | **Error-correction performance** | | |
| --- | --- | --- | --- |
|  | **Mismatches (per 100kb)** | **Indels (per 100kb)** | **Accuracy (%)** |
| Nextera | 0 | 0 | 100 |
| Hybrid |  |  |  |
| Unamplified | 32.13 | 8.02 | 99.97 |
| Virion1 | 25.49 | 6.02 | 99.98 |
| Virion2 | 30.88 | 5.02 | 99.98 |
| Miniasm+R |  |  |  |
| Unamplified | 633.43 | 398.64 | 99.48 |
| Virion1 | 622.37 | 309.42 | 99.53 |
| Virion2 | 519.76 | 249.02 | 99.61 |
| Miniasm+R+M |  |  |  |
| Unamplified | 649.67 | 330.12 | 99.51 |
| Virion1 | 599.39 | 181.3 | 99.61 |
| Virion2 | 518.28 | 152.38 | 99.66 |
| Miniasm+R+M+P |  |  |  |
| Unamplified | 556.69 | 121.56 | 99.66 |
| Virion1 | 607.01 | 84.06 | 99.65 |
| Virion2 | 499.65 | 71.84 | 99.71 |
| Flye |  |  |  |
| Virion1 | 594.26 | 630.14 | 99.39 |
| Virion2 | 560.65 | 519.78 | 99.46 |
| Flye+M |  |  |  |
| Virion1 | 648.38 | 467.83 | 99.44 |
| Virion2 | 613.59 | 368.86 | 99.51 |
| Flye+M+P |  |  |  |
| Virion1 | 622.44 | 147.83 | 99.61 |
| Virion2 | 553.53 | 132.31 | 99.66 |
