## Supplementary Table 3 for "VirION2: a short- and long-read sequencing and informatics workflow to study the genomic diversity of viruses in nature"

**Supplementary Table 3:** Contigs and protein statistics between assembly strategies above 2.5kb

| **Assembly (>2.5kb)** | **General assembly metrics** | | | | |
| --- | --- | --- | --- | --- | --- |
|  | **Nbr. of contigs** | **N50 (bp)** | **Max. contig size (bp)** | **Nbr. of proteins** | **Median protein size (95% CI)** |
| Nextera | 12,977 | 6,193 | 110,882 | 95,642 | 147(146 - 148) |
| Hybrid |  |  |  |  |  |
| Unamplified | 17,342 | 9,042 | 156,874 | 168,453 | 131 (130 - 131) |
| Virion1 | 15,971 | 7,840 | 152,919 | 141,041 | 139 (139 - 140) |
| Virion2 | 16,503 | 8,778 | 152,932 | 155,118 | 137 (136 - 138) |
| Miniasm+R+M+P |  |  |  |  |  |
| Unamplified | 639 | 56,779 | 213,226 | 44,226 | 112 (111 – 113) |
| Virion1 | 834 | 16,840 | 115,168 | 20,398 | 113 (112 – 115) |
| Virion2 | 1,203 | 31,496 | 194,588 | 43,796 | 131 (130 – 132) |
| Flye+M+P |  |  |  |  |  |
| Virion1 | 2,740 | 21,231 | 140,205 | 75,541 | 108 (107 – 109) |
| Virion2 | 4,059 | 31,130 | 570,045 | 153,120 | 110 (109 – 110) |
