## Supplementary Table 4 for "VirION2: a short- and long-read sequencing and informatics workflow to study the genomic diversity of viruses in nature"

**Supplementary Table 4:** Genome-based metrics between Nextera-only and virION-enhanced datasets (>5kb genomes). SD=standard deviation.

|  | **Nextera** | **virION-enhanced** | **virION 2-enhanced** |
| --- | --- | --- | --- |
| Nbr. of viruses | 3,062 | 4,895 | 5,161 |
| Median genome size | 7,972.5 | 9,000 | 9,606 |
| Max. genome size | 110,882 | 152,408 | 194,588 |
| Mean completeness (SD) | 19.51 (17.02113) | 22.4 (20.6592) | 24.44 (22.08735) |
| Mean microdiversity (SD) | 6.939876e-05 (0.0001279526) | 0.0001732822 (0.0003270024) | 0.0001668705 (0.0002889025) |
