## Supplementary Analysis 1 for "VirION2: a short- and long-read sequencing and informatics workflow to study the genomic diversity of viruses in nature"

**Supplementary Analysis 1: Evaluation of the impact of sequencing error on microdiversity estimates**

In this analysis, we evaluated whether SNP frequency analysis of genomes could be influenced by residual errors present on Nanopore-based virus assemblies. In our error-corrected datasets, VirION and VirION 2 OLC assemblies had sequencing accuracies of 99.65% and 99.71%, respectively, with an average of 553 mismatches (per 100kb) remaining post correction (**Supplementary Table 2**). Importantly, validated SNV analysis with short-read corrected Nanopore data has been possible in part due to Nanopore-induced errors being randomly distributed along reads [1], such that repeated SNP patterns, as required for true SNP validation (see Methods), is attainable despite remaining (random) errors. To verify this assumption, we used error-corrected long-read data from our mock community phages (composed of three phage isolates from pure cultures, see Methods), and attempted to detect SNV within these clonal phage populations. Furthermore, this detection was performed at three stages of long-read processing: 1) no correction, 2) Racon, and 3) Pilon (**Supplementary Table 5**). Without correction (i.e., raw Miniasm assembly), 44 SNPs were detected among the contig datasets, and was reduced to 3 SNPs (on a single contig, out of 27) in both Racon- and Pilon- corrected assemblies (in the same contig). This was suggestive that our microdiversity estimates are minimally impacted by remaining sequencing error. Therefore, instead of sequencing error, we reasoned that the higher degree of microdiversity in ‘enhanced’ datasets (compared to short-read assemblies) could stem mostly from the preservation of SNPs within the OLC-derived fraction of the ‘enhanced’ datasets. Indeed, when we looked at the per-genome π value distributions of each of the three constituent assemblies (i.e., hybrid, Spades, and OLC) within VirION- and VirION 2-enhanced datasets (**Supplementary Figure 4)**, we found that in both, the OLC assembly fraction contained viruses with significantly higher degrees of microdiversity compared the hybrid and short-read only assembly fractions (Wilcoxon rank sum test, p*-value* < 2.2 x 10^-16^,). In addition, in both VirION-based datasets there was a significant difference between the hybrid and Spades assemblies (Wilcoxon rank sum test, *p-value* = 0.02 (VirION-enhanced) and *p-value* = 2.852 x 10^-6^ (VirION 2-enhanced)). This suggested that within hybrid assembly alone, in which long-reads are used only for gap closure and solve repetitive regions post De Bruijn graph assembly of the short reads [2], this was sufficient for significant shifts in SNPs frequencies. Although significant, these π value distributions between hybrid and Spades were much more similar to each other compared to the OLC fraction. Taken together, this analysis showed that the OLC-derived fraction of the ‘enhanced’ datasets is the largest contributor to the significant increase in microdiversity in VirION/VirION 2-enhanced viromes, followed by hybrid assembly. In addition, higher microdiversity in ‘enhanced’ datasets compared to short-read data could be explained due to novel viruses uniquely captured with long-reads. However, this is likely not the case, due to our SNP discovery pipeline relying on short read mappings to estimate SNP frequencies along genomes. Therefore, any truly missed viruses by the short-reads, but assembled with long-reads, would not receive a microdiversity estimate due to lack of short-read coverage.
